# Effect of microstimulation of the inferior temporal cortex on color judgment behavior

**DOI:** 10.1101/2023.03.31.535032

**Authors:** Kowa Koida, Hidehiko Komatsu

## Abstract

Within the anterior inferior temporal cortex (AIT) there is a cluster of color selective neurons whose activities correlate with color discrimination behavior. To examine the causal relationship between the activities of these neurons and behavior, we applied electrical microstimulation to modulate neuronal activities within the AIT. We trained monkeys to perform a color judgment task and evaluated the effect of microstimulation in terms of the horizontal shift of the psychometric function. We found large effects of microstimulation on color discrimination behavior, predominantly within a subregion of the AIT. The cortical extent where microstimulation modulated behavior correlated with the presence of color-selective neurons. Unexpectedly, the direction of the modulation of color judgment evoked by microstimulation correlated negatively with the preference of the neurons around the stimulation site. These results support the existence of an anterior inferior temporal color-selective area (AITC) and its causal relationship with the color perception of these animals.

## Introduction

Information about color, an essential and independent attribute of visual stimuli, is transmitted along the ventral visual stream, which includes areas V1, V2 and V4, until it ultimately reaches the inferior temporal cortex (IT) (Conway et al. 2010; Desimone et al. 1985; Fujita et al. 1992; Komatsu 1998; Maunsell and Newsome 1987; Tootell et al. 2004; Zeki 2005). Lesioning or cooling the anterior IT (AIT) severely impairs color discrimination (Buckley et al. 1997; Dean 1979; Heywood et al. 1995; Horel 1994; Huxlin et al. 2000), and imaging and neural recording studies have demonstrated color-selective responses in several subregions of both the AIT and posterior IT cortex (Conway et al. 2007; Conway and Tsao 2006; Harada et al. 2009; Koida and Komatsu 2007; Komatsu and Ideura 1993; Komatsu et al. 1992; Lafer-Sousa and Conway 2013; Takechi et al. 1997). One of these color-selective subregions was detected in the inferior temporal gyrus, near the posterior end of the anterior middle temporal sulcus (AMTS) in the AIT and was named the AIT color area (AITC) (Banno et al. 2010; Namima et al. 2014; Yasuda et al. 2010). Simultaneous recordings of the activities of color-selective neurons in the AITC and color judgment behavior in monkeys has revealed that the activities of these neurons correlate positively with the monkeys’ color judgment (Matsumora et al. 2008). Comparison between the ability of single neurons and monkeys to discriminate colors indicated that variation in the color discrimination threshold across a color space in the monkeys corresponded well with that in the neurons. Although these recording studies suggest a correlation between the activities of color-selective neurons in the AITC and color discrimination behavior, a causal relationship has not yet been demonstrated.

Electrical microstimulation, which enables one to manipulate the activities of small groups of neurons, has been successfully used to demonstrate causal links between neuronal activities in visual cortical areas where specific visual attributes are represented and visual discrimination behavior based on the same attributes (Afraz et al. 2006; Clark et al. 2011; Cohen and Newsome 2004; DeAngelis et al. 1998; Salzman et al. 1990; Verhoef et al. 2012). In the present study, we applied this method to the AITC and measured the effect of microstimulation on color judgment behavior in monkeys. We asked how the magnitude of the effects distribute in the AIT, and whether it is related to the distribution of color-selective neurons (extent of the AITC). In addition we asked whether the effect of microstimulation on color judgment behavior could be explained by the color preference of the neurons at the stimulation site.

## Materials and Methods

### Experimental model

Two Japanese macaque monkeys (*Macaca fuscata*, one female monkey YU weighting 5.5 kg, and one male monkey RG weighing 7.0 kg) participated in the experiments. All procedures for animal care and experimentation were in accordance with the National Institutes of Health Guide for the Care and Use of Laboratory Animals (1996) and were approved by the Institutional Animal Experimentation Committee.

### Preparation of the animals

The surgical procedure was described previously (Matsumora et al. 2008). In short, the animal’s head was fixed with a head holder, and eye position was recorded using an implanted eye coil (1 kHz) (Judge et al. 1980) or a video-based monitoring system (120 Hz, ISCAN, MA, US). The recording chamber was placed on the skull, where an electrode could be inserted vertically into a region of interest in the ventral surface of the IT. This was guided using landmarks in magnetic resonance images obtained before the surgery. Recording positions were identified by comparing the depth profile of neuron activities during each penetration and MRI images, as well as the tip of the electrode identified in x-ray photos. At the end of the experiment, a constant dynamic current was applied to several recording sites to mark the sites and confirm that the electrode tip was located as intended.

### Color judgment task

The monkeys were trained to perform a fine color judgment task (Fig. 1a). A sample color was presented for 500 ms on the fovea while the animal fixated (eye window 2.6 deg) at a distance of 82 cm from the CRT display (CV921X, Totoku, Tokyo, Japan). The fixation spot was turned off 350 ms before the sample onset. The stimulus shape was always a circle with a diameter of 2 deg. The chromaticity of the sample was randomly selected from among 7 colors in one sample set arbitrarily named #1 to #7. Two white targets were presented 5 deg horizontally or vertically from the fixation point just after sample offset. Sample colors #1-3 and #5-7 were associated with different targets. The animals made a saccade to either one of the targets by judging whether the sample color was more similar to either end of the spectrum of 7 colors. A juice reward was given if the monkey made the correct saccade, except for color #4 for which a reward was given randomly (50%). After making a saccade, the color of the two targets changed from white to the colors at either end (#1 and #7) for 100 ms, which facilitated the association between the target direction and sample color. For each recording, the number of repetitions was 20 for monkey YU and 15 for monkey RG, and a total of 280 or 210 trials were performed for each set. The same monkeys participated in an experiment employing similar color judgment tasks for over two years (Matsumora et al. 2008).

**Figure 1.**
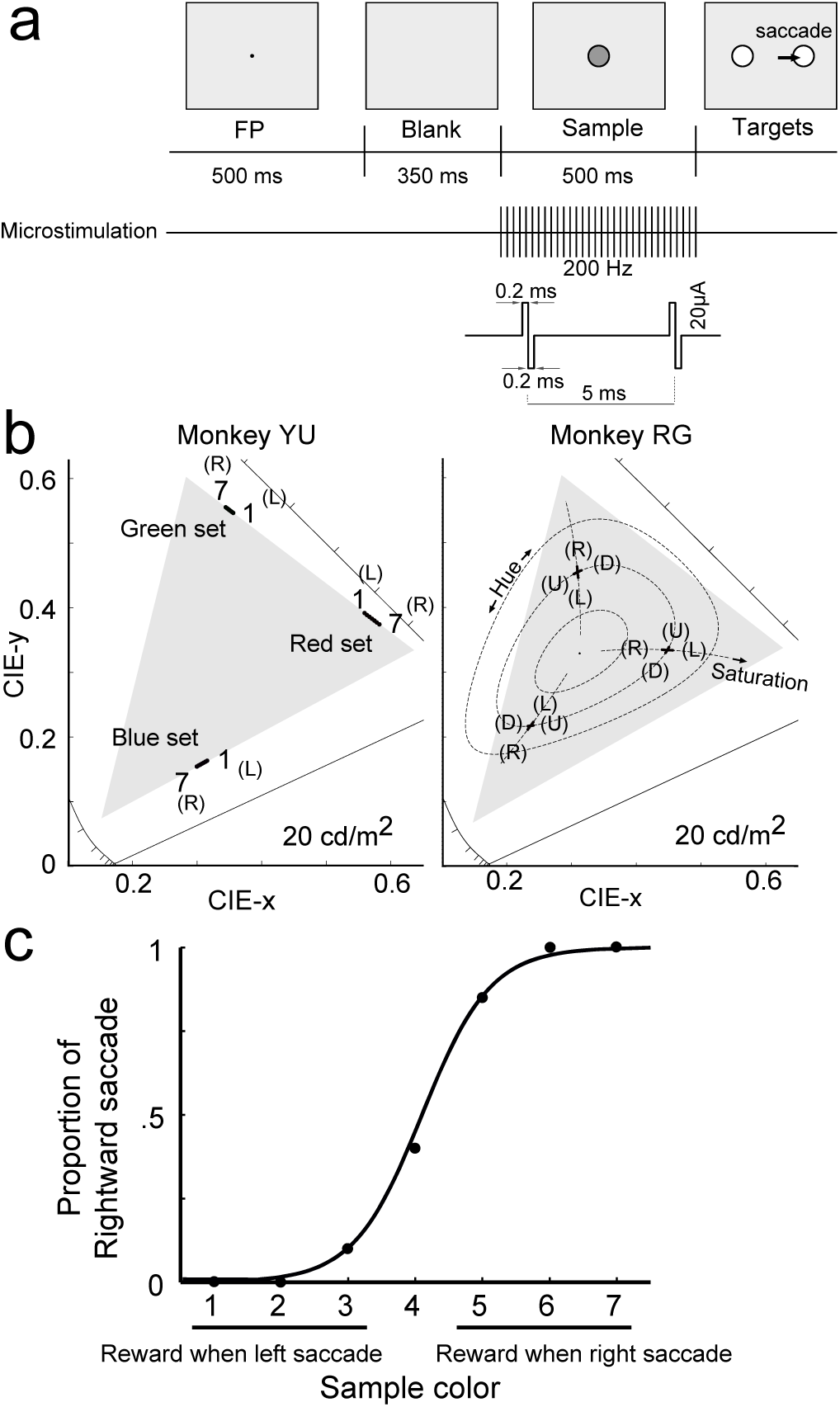
Task design and behavior. (a) Time course of the visual stimulus presentation and electrical microstimulation within one trial. The waveform of the microstimulation pulses is shown below. (b) Chromaticity coordinates of the color sets for monkeys YU (left) and RG (right) shown on a CIE-xy diagram. Three color sets aligned on the edge of the display gamut were used for monkey YU. Six color sets distributed along either the hue or saturation direction were used for monkey RG. Each color set consisted of seven color samples (black dots; overlapped in the figure because of the small color differences). Numbers shown near the black dots indicate arbitrarily assigned sample numbers. Associated saccade directions were left (L), right (R), up (U), or down (D). Gray triangles show the gamut of the display. Dotted lines on the right panel indicate constant hue and saturation loci determined from CIELAB color space. (c) Representative example of the color judgment behavior of monkey YU. The horizontal axis shows the sample color number and associated rules. The vertical axis indicates the proportion of the animal’s binary behavior. Black dots indicate the observed data; the curve is the maximal likelihood logistic function.

### Visual stimulus

Sample color stimuli (20 cd/m^2^) were presented on a gray background (Standard white D65, CIE x = 0.313, CIE y = 0.329, 10 cd/m^2^). The luminance and chromaticity of the stimuli were calibrated using a photometer (CS-200, Konica-minolta, Tokyo, Japan) and presented using a 14-bit linearized video card (VSG 2/3, Cambridge research systems, Rochester, UK). Three sets of sample colors were used for monkey YU and six sets for monkey RG (Fig. 1b).

In our previous study (Matsumora et al. 2008), color stimuli were chosen in every experimental session according to the neuron’s preferred color. In the present study, however, we used three fixed sets of colors for monkey YU and six sets of colors for monkey RG (Fig 1b). We used fixed color sets because 1) microstimulation was applied to sites both with and without color-selective responses and 2) it is not yet clear whether neurons with similar color preferences are clustered within the AIT. Each sample color set consisted of 7 colors. The tests for each color set were separated into blocks.

#### Three edge-color sets for monkey YU

The colors in each of the three sets (green-set, red-set and blue-set) were distributed along the edge of the CRT gamut (Fig. 1b). The seven colors in each set were separated by the same distance on the CIE-xy chromaticity diagram. Color #1 was associated with a leftward saccade and #7 with a rightward saccade. The order of the stimulus blocks was always the same: red, green and blue.

#### Six hue-saturation color set for monkey RG

The seven colors in each of the six sets were distributed in either the hue or saturation direction in the CIELAB color space, a uniform color space. They were separated by the same distance in this color space, and color #4 was common across two color sets that had a hue angle of red (27 deg), green (140 deg) or blue (−70 deg) and a saturation of 40 unit radius from the neutral point. The constant saturation locus is circular in this color space, but we approximated it as linear because of the small color distance. Behavioral responses were a rightward or leftward saccade for the saturation color set (rightward saccades associated with desaturated red, saturated green, and saturated blue), and upward or downward saccades for the hue color sets (upward saccades associated with yellowish red, bluish green, and reddish blue, Fig 1b). This rotation of saccade direction was introduced to avoid crosstalk between the behavioral responses to hue and saturation. For example if microstimulation modulates color perception toward both hue and saturation, the animal would judge color in the hue or saturation directions independently. The order of the stimulus blocks was red-hue, green-hue, blue-hue, red-saturation, green-saturation and blue-saturation.

The range of colors between #1 and #7 in each color set was determined after extensive training of the monkeys in a color judgment task, when their behavioral performance had arrived at a plateau level. The interval of the color set was determined such that both extremes of the behavioral responses were clearly observed (0 or 100%), and the intermediate response ratio was observed with several stimuli (Fig. 1c).

When the stimulus block was changed, several trials under both the non-stimulation and stimulation conditions were performed as brief training. These data were not used for analysis. The influence of the order of the color sets was tested by repeating the experiment with different initial color sets, and we confirmed that the stimulation effect was reproducible.

### Electrical Microstimulation

For microstimulation, in half of the trials biphasic electrical pulses (200 Hz, anode first; Fig. 1a) were randomly applied through the same electrode used for neural recording (0.5-1.5 MΩ, FHC, ME, US). The stimulus current was usually 20 μA, but was 50 μA for the right hemisphere of monkey RG. The duration and timing of the stimulation were the same as for the visual stimuli (500 ms). The reward rule was the same with or without stimulation.

### Penetration and recording

Neural recordings were made using electrodes inserted through an evenly spaced grid of holes at 1-mm intervals (Crist et al. 1988) over a wide area of the ventral surface of the IT. Electrodes were penetrated vertically and entered the target cortex from a deeper layer. The experiment was performed when the electrode was advanced more than 200 μm from the entry depth of the deep layer. In one electrode penetration, we performed the experiments up to three times, and the electrode was displaced at least 300 μm in each experiment. At each site, we performed unit recording and microstimulation through the same electrode. We first examined the color selectivity of the multi-unit activity (MUA) elicited by 15 equiluminant color stimuli that evenly distributed across the color gamut on the CIE-xy chromaticity diagram while the animals were fixating. Then the effects of microstimulation on color judgment behavior were measured. Neural activities were amplified, filtered and sampled at 25 kHz, and MUAs were detected by a setting threshold such that the mean firing rate at baseline (300-ms period before sample onset) became 40 spikes per second. Net responses were calculated after subtracting the baseline activity and then used for the analysis.

### Stimuli for the fixation task

To examine the color selectivity of neuronal responses, we used 15 equiluminant (20 cd/m^2^) color samples arranged systematically on the chromaticity coordinates (Fig. 2b, d), except for saturated blue (10 cd/m^2^). The stimulus shape for the color-selectivity test was a circle in most cases, but other geometrical shapes were used in some cases when the response to that shape was significantly larger than to a circle.

**Figure 2.**
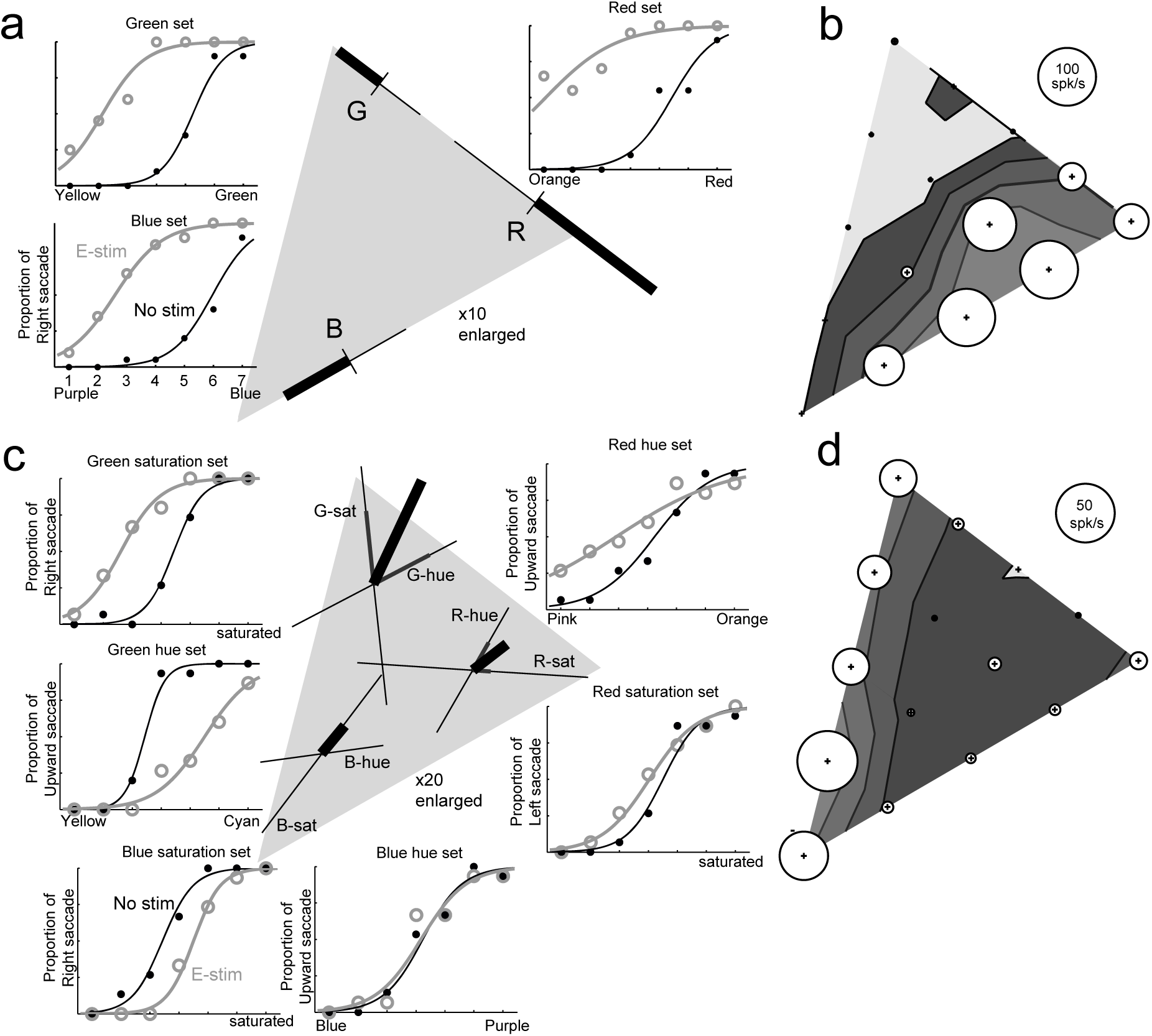
Effect of microstimulation and color-selective responses. (a) Example results from monkey YU. Three inset panels show the animal’s behavior in the stimulation (circles with gray line) and non-stimulation (dots with black line) conditions. The panels depict results obtained with the red, green and blue sets. Fittings were done using two independent logistic functions. The center panel summarizes the effects of microstimulation defined as the horizontal shift of the functions shown on the chromaticity diagram (black bars). Note that the length of the bar was magnified 10 times for the purpose of visualization. The filled gray triangle indicates the display gamut in CIE-xy color space. Thin lines show the sample color range from #1 to #7 (also magnified 10 times), and orthogonal ticks indicate the position of sample #4. (b) Color-selective multiunit activity recorded from the same electrode position used for the microstimulation experiment in (a). Circle diameters indicate response magnitudes to each of 15 color samples recorded while the animals performed a fixation task. Linearly interpolated response contours are also shown. (c) Example results from monkey RG. Six inset panels show the results with six color sets. Symbols are the same as in (a). The center panel summarizes the effects of microstimulation with the six color sets. Crossed thin lines indicate the ranges of the hue and saturation sets (magnified 20 times). Gray bars on the thin lines indicate the horizontal shift for each color set, and thick bars indicate their vector sum. The lengths of the bars were magnified 20 times for visualization purpose. (d) Color-selective multiunit activity recorded from the same electrode position used for the microstimulation experiment shown in (c); the format is same as in (b).

### Analysis of behavior

Behavioral performance was fitted by a maximum likelihood logistic equation using a generalized linear model (glmfit function in Matlab). The observed psychometric function was characterized by its horizontal position and slope at the point corresponding to 50% of the height of the function. The effect of microstimulation was summarized as the horizontal shift and slope change between two independently fitted curves; values were obtained by subtracting data acquired in the non-stimulation condition from data acquired in the stimulation condition. The significances of the horizontal shift and slope change were tested using a permutation test, where the animals’ responses under both stimulation and non-stimulation conditions were mixed, then randomly separated into two conditions while keeping the visual stimulus condition constant. The horizontal shift and slope change were calculated by fitting the resulting data. This process was repeated 2,000 times and p values were calculated based on the distributions of each variable. The horizontal shift and slope change were first quantified on the stimulus metrics (#1 to #7), then were normalized to mean behavioral performance to allow comparison across different color sets and animals. One unit of horizontal shift corresponded to a change in performance from 50% to 86% in the logistic function, which corresponded to the average standard deviation (SD) of the function.

In two experimental blocks for monkey YU, the psychometric function in the stimulating condition exceeded the range of the color set because of the large effect of microstimulation. In those cases, data were clipped at 5 SD units for further analysis (Fig. 3).

**Figure 3.**
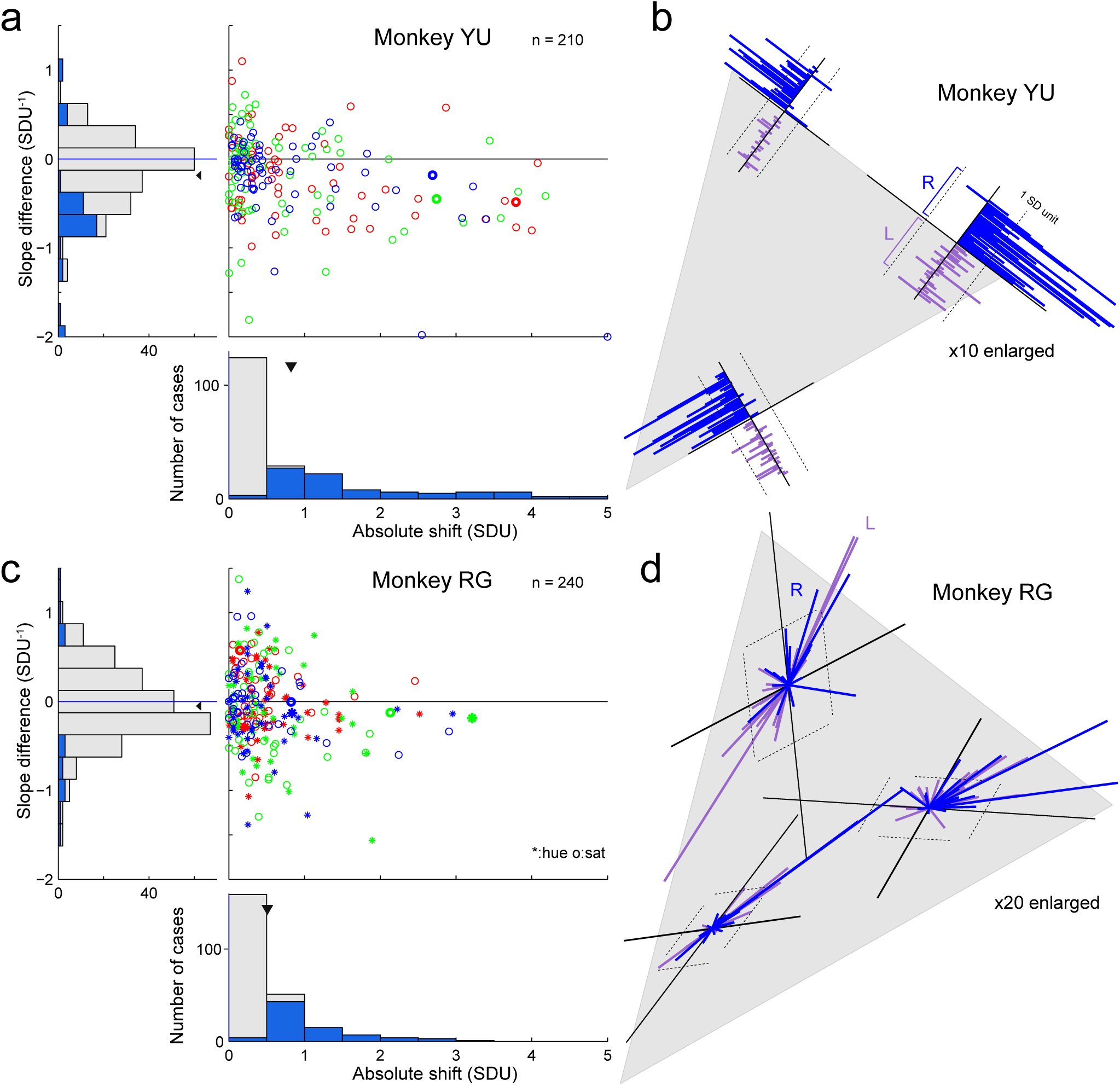
Summary of the effects in all experiments. (a) Effects of microstimulation in monkey YU summarized as an absolute horizontal shift (horizontal axis) and slope change (vertical axis). Histograms for each axis are shown to the bottom and left; the colored region indicates the cases significantly different from zero (p < 0.05, permutation test). Triangles above each histogram indicate the mean. Colors of the circles on the scatter diagram indicate the color set. Thick circles correspond to example data in Figure 2a. (b) Direction and size of the shifts for monkey YU shown in CIE-xy color space; the data were magnified 10 times for visualization purposes. The origin of each bar plots corresponds to sample color #4, and each bar represents data from an individual experiment. Bars are separated for each hemisphere (L and R denote data from the left and right hemispheres) and arranged according to the order of the experiment. Dotted lines indicate the mean color judgment performance (1 SD unit). The gray triangle represents the display gamut. (c) Effects of microstimulation in monkey RG. Format is the same as in (a), except asterisks and circles in the scatter diagram represent data for the hue and saturation sets, respectively. (d) Effects of microstimulation in monkey RG shown as vector sums of the horizontal shifts across the hue and saturation color sets. The data were magnified 20 times for visual purposes. Solid black lines indicate the axis of sample color sets. Other conventions are the same as in (b). See also Figures S1-S3.

## Results

Stimulated sites were systematically distributed on the ventral surface of the AIT. The sites were centered at the posterior end of the AMTS, where many color-selective cells are concentrated (Banno et al. 2010; Koida and Komatsu 2007; Komatsu and Ideura 1993; Komatsu et al. 1992; Matsumora et al. 2008). For each of the 76 penetrations, the experiment was performed at up to three sites at different depths. In total, 110 experiments were performed (70 with monkey YU, 40 with monkey RG, Table 1).

**Table. 1.** Number of microstimulation sites and observed significance “Tested” indicates the number of microstimulation sites. Three (monkey YU) or six (monkey RG) color sets were used for each stimulation site. For horizontal shift, “Any” indicates the number of sites where a significant horizontal shift of the psychometric function was observed with at least one color set (p < 0.05). In parentheses are the numbers of sites where significant horizontal shifts of the psychometric function (p < 0.05) were observed with the red, green or blue set in monkey YR, and with the red hue (R-hue), red saturation (R-sat), green hue (G-hue), green saturation (G-sat), blue hue (B-hue) or blue saturation (B-sat) in monkey RG. For slope change, “Any” indicates the number of sites where a significant slope change in the psychometric function (p < 0.05) was observed for at least one color set. Because significant changes in the horizontal shift were observed with more than one color set at multiple sites, the sum of the numbers of cases showing significant changes exceeds the number of sites.

| Monkey YU |  | Horizontal shift |  |  |  | Slope change |
| --- | --- | --- | --- | --- | --- | --- |
| Hemisphere | Tested | Any | Red | Green | Blue | Any |
| R | 34 | 33 | 28 | 22 | 22 | 19 |
| L | 36 | 12 | 5 | 4 | 6 | 5 |
| Both | 70 | 45 | 33 | 26 | 28 | 4 |

| Monkey RG |  | Horizontal shift |  |  |  |  |  |  | Slope change |
| --- | --- | --- | --- | --- | --- | --- | --- | --- | --- |
| Hemisphere | Tested | Any | R-hue | R-sat | G-hue | G-sat | B-hue | B-sat | Any |
| R | 16 | 10 | 5 | 2 | 4 | 3 | 2 | 2 | 4 |
| L | 24 | 19 | 14 | 8 | 13 | 7 | 9 | 8 | 10 |
| Both | 40 | 29 | 19 | 10 | 17 | 10 | 11 | 10 | 14 |

### Large impact on color judgment

We found that microstimulation modulated color-judgment behavior at some stimulation sites. Figure 2 shows the effects of stimulation at two sites where clear modulation of color-judgment behavior was observed. In an example from monkey YU (Fig. 2a, b), the MUA at the stimulation site indicated a preference for pink (right-bottom area in the color space, Fig. 2b). In this case, the effect of microstimulation was apparent in all three color sets, and the psychometric function shifted horizontally when microstimulation was applied. The leftward shift in the psychometric function for the green set indicates the monkey more frequently judged the sample colors as greenish when the microstimulation was applied. Likewise, leftward shifts for red and blue sets indicate the animal more frequently judged the sample colors as reddish and bluish, respectively.

The magnitude of the effect was quantified through logistic regression of the psychometric functions (see Method). Each function was described by two parameters: the horizontal position and the slope. The horizontal position indicates the sample number where the animal chose the two alternatives with equal frequency. The slope indicates the accuracy of the animal’s behavior. Changes in the psychometric function caused by microstimulation were quantified based on the horizontal shift and slope change between conditions with and without stimulation. If microstimulation induced a color signal, a horizontal shift would be expected. If microstimulation induced noise and disturbed choice behavior, irrespective of color, a decrease in slope would be expected. Horizontal shifts caused by stimulation in the example case shown in Figure 2a were −3.13, −4.37 and −3.37 for green, red and blue sets, respectively, which significantly deviated from zero (p < 0.05, permutation test). Slope changes caused by stimulation were −0.513, −0.558 and −0.229 for the green, red and blue sets, respectively.

To evaluate the effect of microstimulation in terms of perceptual biases in a color space, we converted the magnitudes of the horizontal shifts of the psychometric functions into distances in chromaticity (bar length in Fig. 2a). In these examples, microstimulation biased color judgment toward more reddish colors for the red set, more greenish colors for the green set and more bluish colors for the blue set. In all three cases, color judgment was biased in the outward direction in the color space; in other words, toward more saturated colors.

Microstimulation at some sites also modulated the behavior of monkey RG, and an example case is shown in Figure 2c. In this case, the sizes of the effects differed depending on the color set: large effects were observed for four color stimulus sets (red-hue, green-hue, green-saturation and blue-saturation sets, p < 0.05, permutation test), while the effect for the other two color sets was small (red-saturation set) or absent (blue-hue set). To visualize the effect for each color set, the magnitude of the horizontal shift in the psychometric function was converted to distance in chromaticity and was plotted along the direction of each of the hue and saturation color sets on the chromaticity diagram (two thin lines crossing at the center of the chromaticity of each color set in Fig. 2c). Then the effects of two components, in the hue and saturation directions, were combined to make a single vector for the red, green and blue regions (thick lines in Fig. 2c). On the whole, the observed effects tended to distribute toward the upper right in the color space, which is headed away from the blue region. The MUA recorded at this site showed clear color selectivity and preferred blue and green colors (Fig. 2d).

Not only did microstimulation alter the animals’ behavior in the stimulated condition, it also affected behavior in the non-stimulated condition. For example, in the data from monkey YU described above, the psychometric function shifted leftward in the stimulation condition, but rightward in the non-stimulation condition (Fig. 2a). This contrasted with the fact that the psychometric function was centered at sample color #4 for the training session (Fig. 1c) or for the experimental session when microstimulation had no effect (blue-hue set in Fig. 2c). The shift observed in the non-stimulation condition is likely due to *’probability matching’* behavior, where monkeys made a roughly equal number of responses toward both directions over the course of an experiment, and similar effects were reported previously as *’null choice bias’* (Salzman et al. 1992; Verhoef et al. 2012).

### Population results

Across the experiments at the 110 sites tested, significant horizontal shifts of the psychometric function (p < 0.05, permutation test) for at least one color set were observed at 74 sites (45 for monkey YU and 29 for monkey RG). Significant changes in the slope for at least one color set were observed at 38 sites (24 for monkey YU and 14 for monkey RG). A complete list of the numbers of significant cases is summarized in Table 1. The magnitudes of the effects evaluated in terms of the absolute horizontal shift and slope change for all stimulation sites are summarized in Figure 3a and c. Each value was normalized to the mean accuracy of the animal’s judgment for each color set (standard deviation (SD) of the psychometric function) to enable comparison among different color sets and between the animals (see Method). The magnitudes of the horizontal shifts were widely distributed up to 5 SD units (data were clipped when magnitude of the shift exceeded 5 SD units in two cases for monkey YU). Significant horizontal shifts (p < 0.05, permutation test) were observed for 164 color sets (87 for monkey YU, 77 for monkey RG). These cases were not limited to specific color sets (33, 26 and 28 cases for red, green and blue sets for monkey YU and 19, 10, 17, 10, 11 and 10 cases for red-hue, red-saturation, green-hue, green-saturation, blue-hue, blue-saturation sets for monkey RG, respectively). Mean absolute horizontal shifts were 0.82 SD units for monkey YU and 0.43 SD units for monkey RG. When the shifts along two axes (hue and saturation directions) were integrated in Euclidean distance for monkey RG, the average shift was 0.67 for all color sets, and 0.65, 0.83 and 0.52 for red, green and blue integrated sets, respectively (Fig. S1). When the effective sites were considered, the mean absolute horizontal shifts for the different color sets were comparable (p > 0.10, 1-way ANOVA), indicating that the effect of microstimulation was not limited to a specific color set.

The magnitude of the slope change distributed around zero, but its mean was slightly lower than zero (Fig. 3a, c) (−0.19 for monkey YU, −0.05 for monkey RG). Significant changes in the slope for each experiment were observed in 58 cases (42 for monkey YU, 16 for monkey RG, p < 0.05, permutation test), among which negative changes in slope were more frequently observed (negative n = 36, positive n = 6 for monkey YU, negative n = 10, positive n = 6 for monkey RG). Negative changes in the slope indicate microstimulation decreased the stability of the judgments.

There was weak but significant correlation between the horizontal shift and the slope change; the correlation coefficients were −0.40 for monkey YU (p < 0.01, t-test) and −0.17 for monkey RG (p < 0.01, t-test). The negative correlation indicates that when microstimulation biased a monkey’s judgment, it also decreased the stability of the judgment. Because significant horizontal shifts were more frequently observed than significant slope changes for both monkeys, in the following analyses we will focus on the horizontal shift as an index of the effect of microstimulation.

### Direction of modulation in a color space

We next examined the direction of the modulation of the population data, and asked how those directions distribute in a color space. The directions of the shifts for each color set in each experiment summarized in the color space are shown in Fig. 3b, d. As was illustrated in the example case in Fig. 2a, there was a consistent tendency for microstimulation to modulate monkey YU’s judgment toward more saturated colors or toward the corner of the color triangle in the display gamut. This tendency was particularly obvious when the microstimulation was applied to the right hemisphere (blue bars), but it was significant for both hemispheres (p < 0.05, t-test). This suggests stimulation may have made the stimulus color appear more saturated.

To examine this possibility, in monkey RG we used different color sets, one laid along saturation directions and the other along hue directions. However, we did not observe consistent shifts along the saturation direction. Instead the induced shift distributed in various directions, and we observed that large effects were frequently observed along a line from blue or toward blue (Fig. 3d). To quantitatively examine whether there was bias in the effects of microstimulation, we calculated the covariance of the horizontal shifts between the hue and saturation axes. Significant covariance was observed between the hue and saturation sets (correlation coefficient r = 0.55, −0.75, −0.89 for red, green and blue sets, respectively, p < 0.01 for all cases, t-test), which indicates that the directions of the modulation in a color space are biased in certain directions. When extrapolating the directions of modulation for all color sets, they approximately converged at a blue region in the left bottom of the chromaticity diagram. When this extrapolation was performed in cone excitation space, which is another coordinate set of color stimuli based on photoreceptor outputs, the directions of modulation did not align along the S cone axis, but tended to rotate clockwise (Fig. S2). We conclude that there is a consistent tendency in the direction of modulation in the color space.

### Mapping cortical topography

To examine the spatial relationships between sites where the effects of the microstimulation were observed and where color-selective neural responses were recorded, we compared their distributions on the ventral surface of the IT cortex (Fig. 4). Hemispheric bias in the effect of microstimulation was observed in both monkeys: larger effects were frequently observed in the right hemisphere of monkey YU and the left hemisphere of monkey RG (Fig. 4c, d). This bias was not due to the order of the experiments, as the experiments with monkey YU were not conducted serially but were switched several times between hemispheres. Strong impacts were observed in a subregion extending a few millimeters and positioned adjacent to the posterior end of AMTS. This localization was again not explained by the experimental order, because the penetration sites were pseudorandomly chosen within each hemisphere.

**Figure 4.**
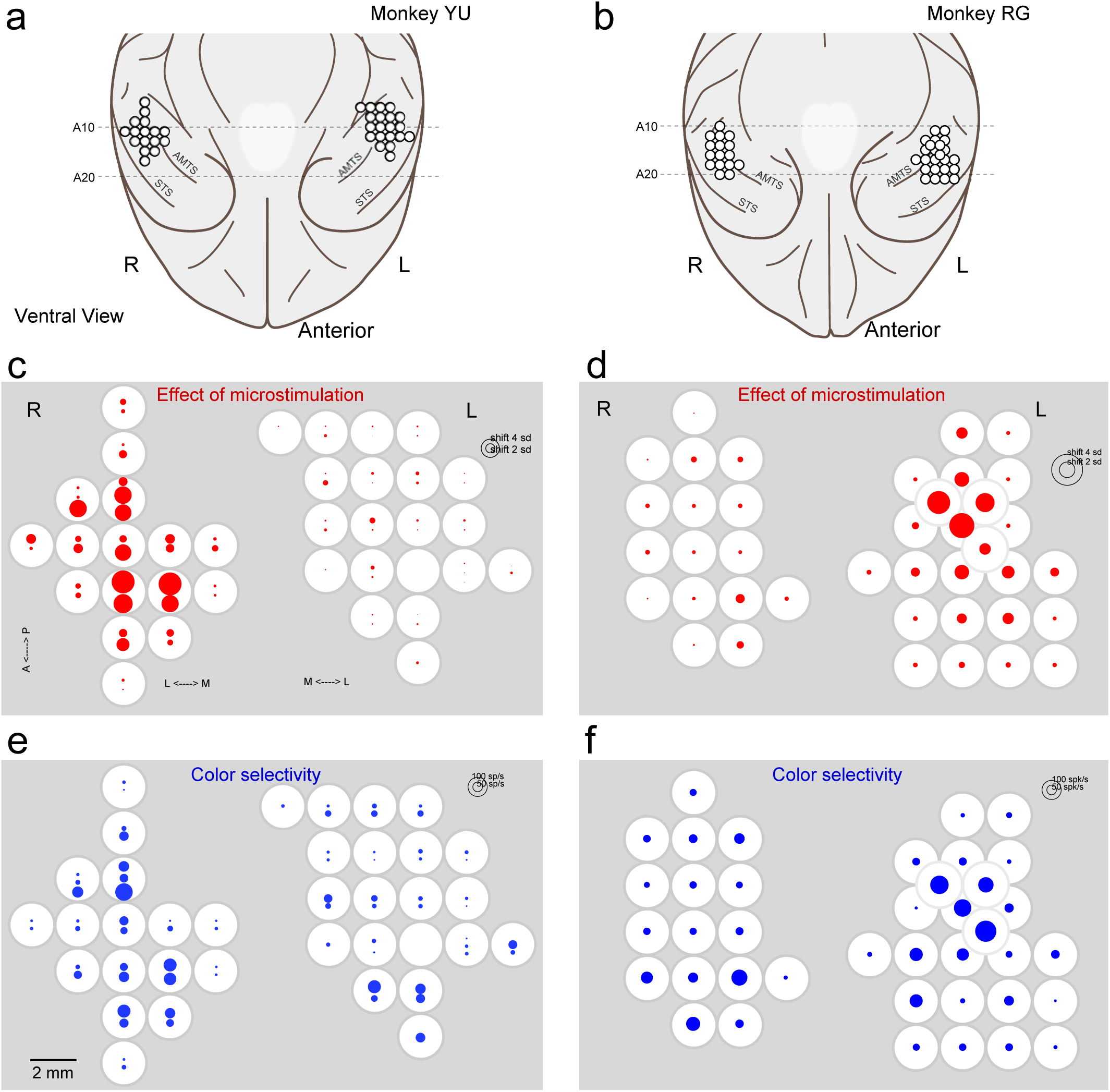
Distribution of the effect of microstimulation and color-selective activities. (a, b) Ventral views of the anterior parts of the cerebral cortex in monkeys YU (a) and RG (b). Sites of neural recording and stimulation are shown by open circles. The intervals between penetrations were usually 2 mm. Dashed lines indicate 10 and 20 mm anterior from the interaural line (A10 and A20, respectively). AMTS, anterior middle temporal sulcus. STS, superior temporal sulcus. (c, d) Map for the effect of microstimulation in each hemisphere. Circle diameters indicate the sizes of the horizontal shifts elicited by the stimulations. Three to six color sets were tested at each site, and the maximum horizontal shift among them is shown. White circles indicate penetration sites and correspond to those in (a). In monkey YU, two or three stimulations were conducted at different depths, and the results are shown separately within the white circles: upper symbols represent the data from deeper layers. (e, f) Map of the color-selective responses of multiunit activities recorded at the same electrode position as in (c, d). Diameters of blue circles indicate the difference between maximum and minimum responses among 15 equiluminant color stimuli.

Color-selective responses in the MUA were also clustered within the IT region examined (Fig. 4e, f). The strength of the color selectivity was assessed as the difference between the maximum and minimum responses across 15 equiluminant colors. We also performed similar analyses using the maximum response or the difference between the maximum and minimum responses divided by the maximum as a measure of the strength of the color selectivity, and the results were similar. Strong color-selective responses were observed in both hemispheres in both monkeys. Clusters of color-selective sites were observed in the right hemisphere of monkey YU and left hemisphere of monkey RG, where they extended several millimeters adjacent to the posterior end of the AMTS, which is consistent with our earlier findings (Banno et al. 2010; Koida and Komatsu 2007; Komatsu and Ideura 1993; Komatsu et al. 1992; Matsumora et al. 2008).

To assess the coincidence of the spatial distribution between the effect of microstimulation and color-selective responses, we examined their correlation (Fig. 5). A significant correlation was observed for both monkeys when the data from both hemispheres were pooled (r = 0.564 for monkey YU, r = 0.559 for monkey RG, p < 0.05, t-test). When the data from each hemisphere was analyzed separately, a significant correlation was observed for the effective hemisphere (r = 0.670 for the right hemisphere of monkey YU, r = 0.665 for the left hemisphere of monkey RG, p < 0.05), but not for the other hemispheres. Significant correlations indicate that the spatial organization of the effect of microstimulation coincided with that of the color-selective MUA.

**Figure 5.**
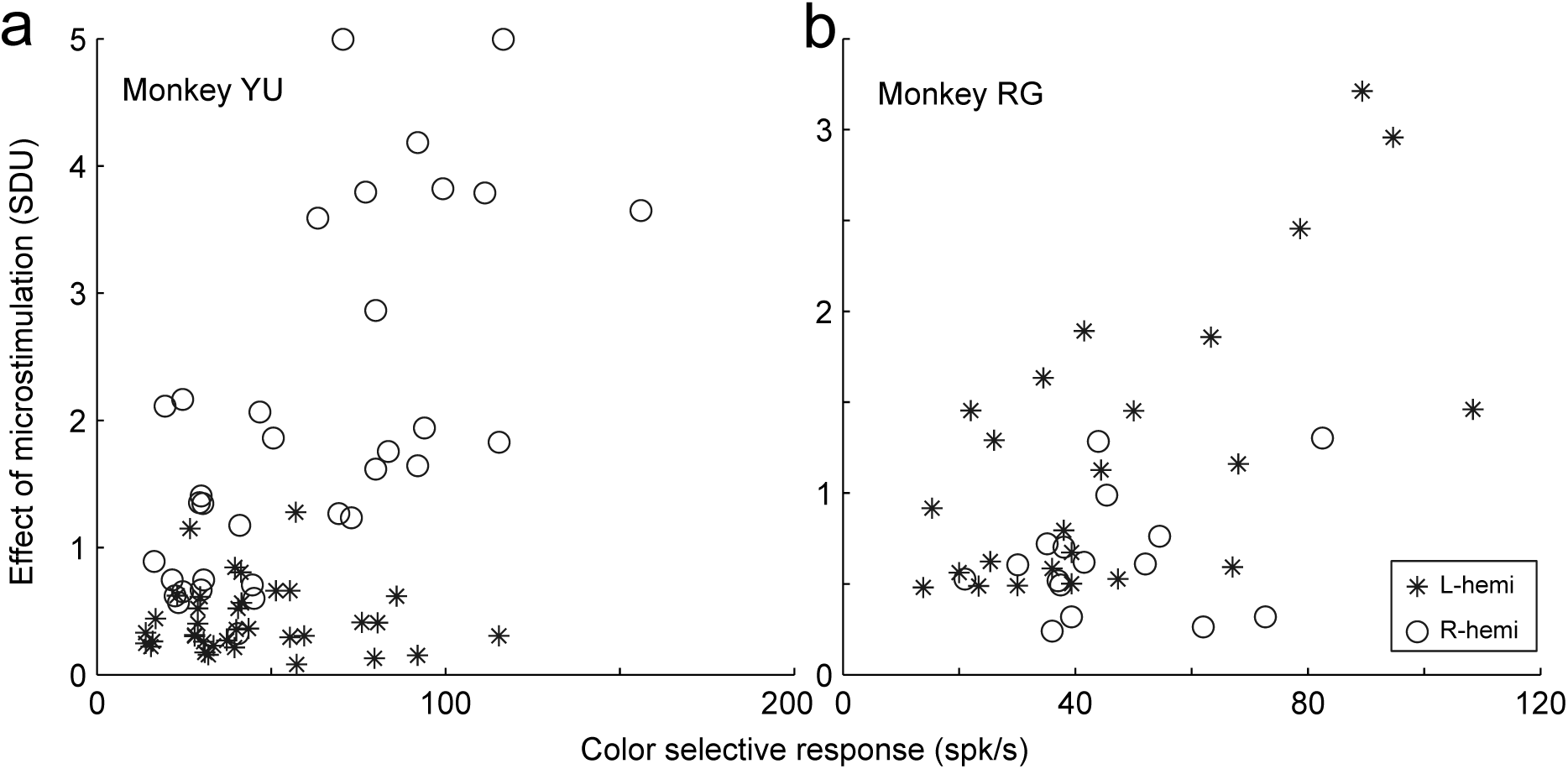
Relationship between the effect of microstimulation and neural color selectivity at the same site in monkeys YU (a) and RG (b). Circles and asterisks indicate the data from the right and left hemispheres, respectively. The horizontal axis represents color selectivity and is the same as in Figure 4e, f. The vertical axis represents the effect of microstimulation in SD units and is the same as in Figure 4c, d.

### Comparison between direction of modulation and color preference

We next examined whether there is any relationship between the preferred color indicated by the MUA at each recording site and the direction of modulation in a color space. Analysis was performed only in the right hemisphere of monkey YU and the left hemisphere of monkey RG, where effects of microstimulation were clearly observed.

Three example cases recorded from different penetration sites in monkey YU are shown in Figure 6a-c. Among 15 sample colors, the preferred colors indicated by the maximum MUA responses were pink (bottom right in the color space, Fig. 6a), orange (upper right, Fig. 6b), and red (upper right, Fig. 6c). The directions of modulation in color judgment evoked by microstimulation (blue bars) were compared with color preference vectors (orange bars) that connected from the stimulus color to the preferred colors. For the MUA depicted in Fig. 6a, the directions of the color preference vectors were downward for the green set, down-leftward for the red set, and up-rightward for the blue set (Fig. 6a). On the other hand, the directions of the biases evoked by microstimulation all pointed toward the corner of the gamut. These two sets of directions tended to be in the opposite or orthogonal directions with respect to each other. Similar tendencies were also observed in the other example cases (Fig. 6b, c). Because the direction of the bias elicited by microstimulation in monkey YU was estimated from only a one-dimensional color judgment test, the direction of the bias in a two-dimensional color space could not be determined.

**Figure 6.**
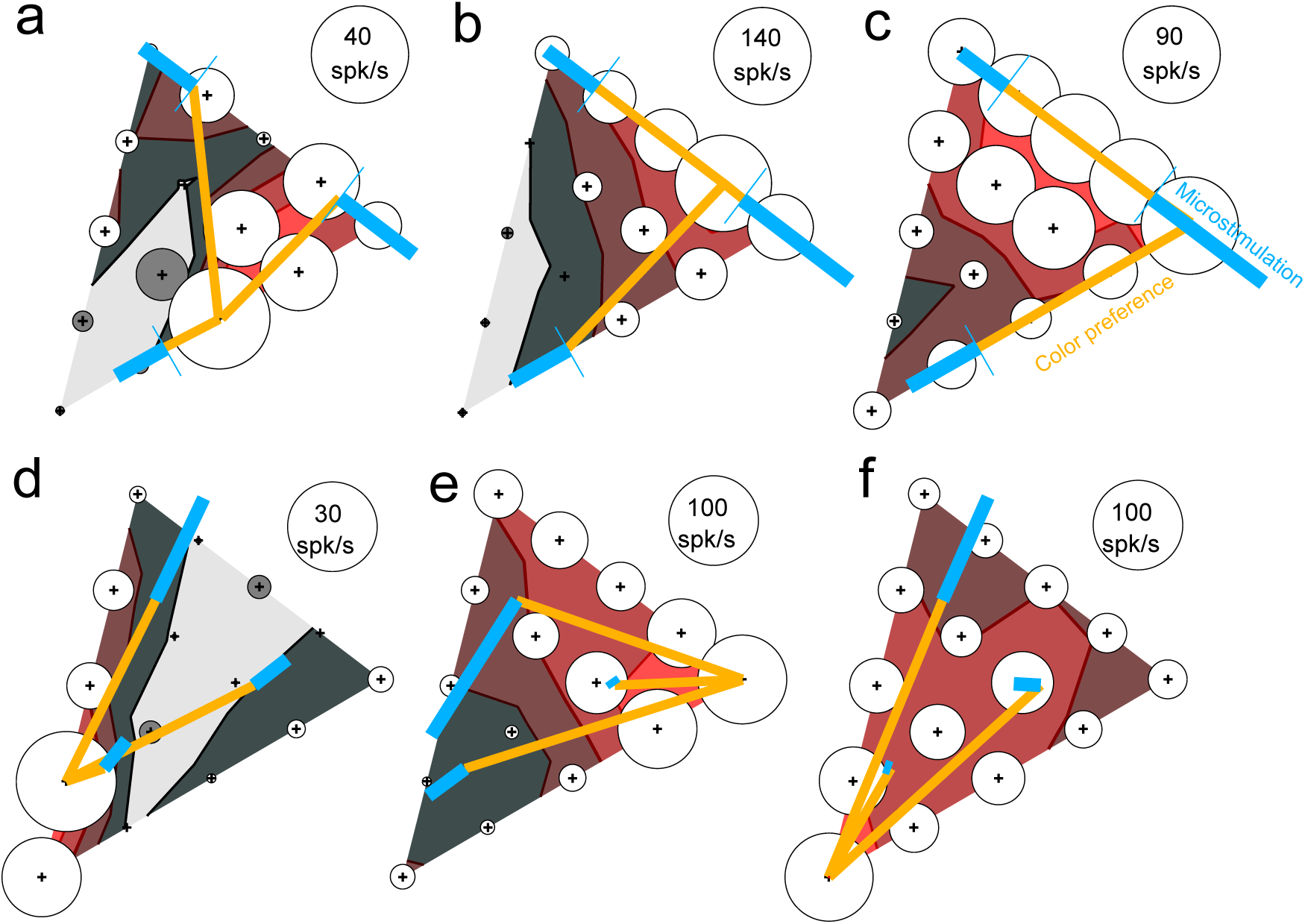
Comparison between the direction of the effect of microstimulation and the color selectivity at three example sites in monkeys YU (a-c) and RG (d-f). Effects of microstimulation are superimposed on the color selectivity plot for each site (bubble and contour plot as shown in Fig. 2). Blue bars indicate the direction of the effect of microstimulation; the format is the same as in Figure 3. Orange bars connect the chromaticity coordinates of color #4 in the sample color set of the color judgment task and the preferred color among the 15 colors used to test color selectivity. The example case in (d) is the same as depicted in Figure 2c.

For monkey RG, we estimated the direction of the bias in two dimensional color space based on the two experiments using the hue set and saturation set for each color region. Thus comparison between the direction of the behavioral shift and color preference indicated by the MUA would be more straightforward than that for monkey YU. In the first example (Fig. 6d), microstimulation biased color judgment toward top-right for all three color sets, which is away from the preferred color (blue, bottom-left). This means the shift and color selectivity were in opposite directions. In the second example (Fig. 6e), the directions of the bias tended to converge toward the blue color (down left direction), and the preferred color indicated by the MUA (red, right corner) was roughly in the opposite direction. In the third example, a clear shift was observed only for the green sample set (Fig. 6f), and it was directed toward the yellow-green color. The preferred color indicated by the MUA at this site was blue, though the selectivity was rather broad. Nonetheless, the shift for the green set was again in the direction opposite to the preferred color.

To quantitatively compare the directions of the modulation induced by the microstimulation and the direction of the preferred color indicated by the MUA at the same site, we computed the angle difference between these two directions (Fig. 7). Although the observed directions were widely distributed, there was clear bias toward the opposite half with both monkeys. If we consider only the cases showing large effects of microstimulation (bias > 2.5 SD), for monkey YU, there were 4 cases with a similar direction (< 90 deg) and 17 cases with an opposite direction (> 90 deg, Fig. 7a). This ratio was significantly different from chance (p < 0.01, sign test). For monkey RG, there was only one case with a similar direction but 6 cases with an opposite direction (Fig. 7b). This tendency did not change if a different threshold on the effect of microstimulation was applied. The mean angle difference was 119 deg for monkey YU and 131 deg for RG, both of which were significantly displaced from 90 deg (p < 0.01, t-test). These results indicate that the direction of modulation by the microstimulation tended to lie in the direction opposite to that of the preferred color at the same site.

**Figure 7.**
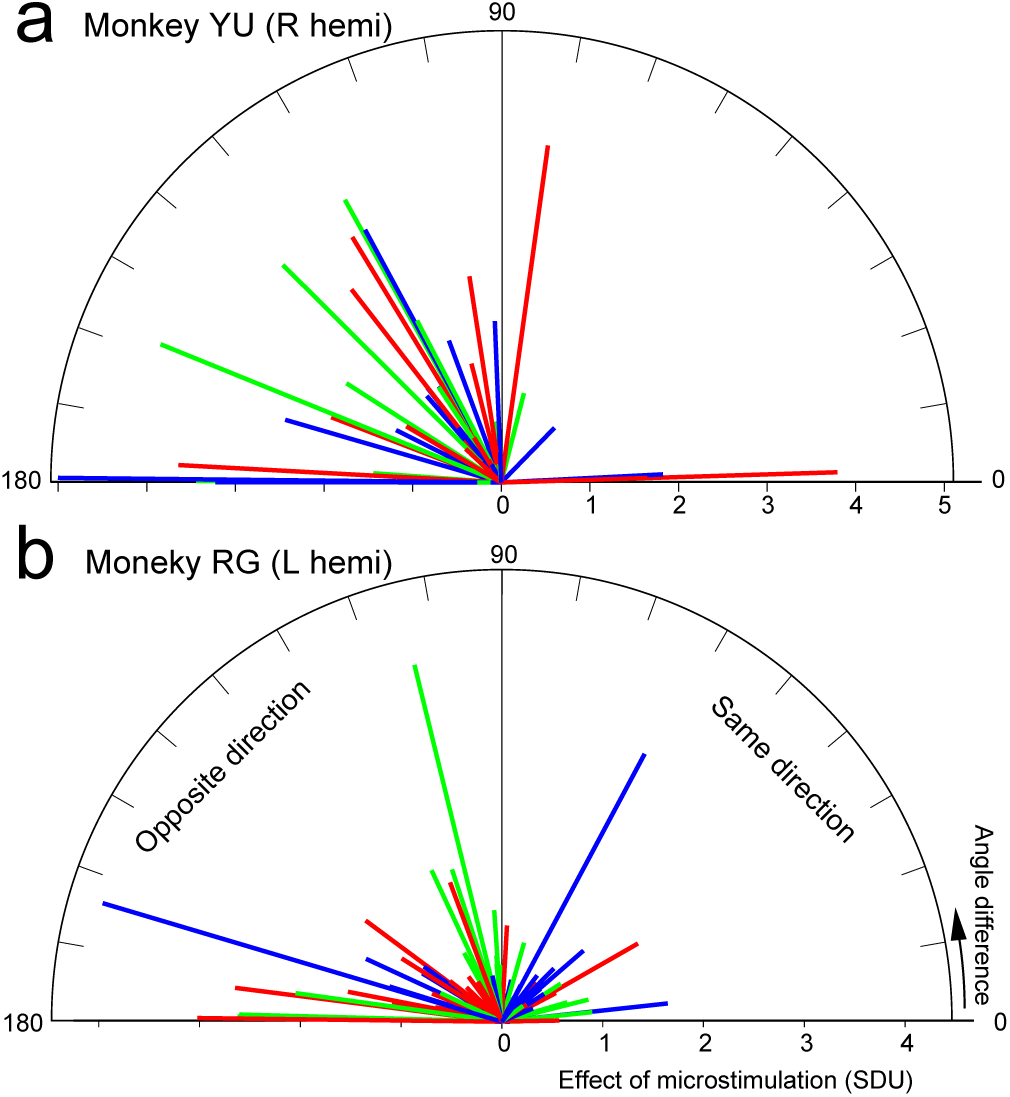
Angle difference between the direction of the modulation and the preferred color for monkeys YU (a) and RG (b). Bar direction indicates absolute angle difference between them, and bar length represents magnitude of the horizontal shift caused by the microstimulation. Bars in the right half (0-90 deg) indicate that the horizontal shift and preferred color are in the same direction; bars in the left half (90-180 deg) indicate they are in opposite directions. Bar color indicates the color set.

## Discussion

We found that microstimulation in the AIT significantly modulated color-judgment behavior. The sites where stimulation modulated the monkeys’ behavior were localized within a subregion of the AIT, and effective sites coincided with the presence of color-selective neurons. The effect of microstimulation was mainly observed as a horizontal shift in the psychometric function and was sometimes accompanied by a decrease in the slope of the function. The horizontal shift in the psychometric function indicates that microstimulation biased the monkeys’ color judgment, and the direction of the bias was toward more saturated colors in monkey YU and away from blue in monkey RG. The bias also tended to be in the direction opposite to the preferred color indicated by the MUA at each stimulation site.

### Potential of factors other than color to induce a stimulation effect

The IT plays important roles in shape and object recognition (Desimone et al. 1984; Logothetis and Sheinberg 1996; Perrett et al. 1982; Rolls 1984; Tanaka 1996) and in color discrimination. Microstimulation may have thus modulated shape or object perception, though this is unlikely. If microstimulation had modulated shape or object perception, it would not have provided any cue affecting color judgment, and no consistent bias in color judgment would be expected.

Likewise, spatial bias due to microstimulation affecting saccade eye movements is unlikely the cause of the observed modulation. Although the receptive fields of cells in the AIT cover both visual fields, there are contralateral biases, and microstimulation may bias judgments in contralateral directions. However, this is not consistent with the observed direction of the bias. For monkey YU, the observed biases across the population tended to be rightward, in the direction of saturated color, regardless of the stimulated hemisphere (Fig. 1b). Furthermore, large effects were dominantly observed in the right hemisphere, and an increased frequency of rightward saccade is hard to reconcile with the biased activities of neurons in the right hemisphere. With monkey RG, we observed no consistent biases toward either leftward or rightward saccade responses with the saturation sets (Fig. 3d). Moreover, the behavioral responses of monkey RG to the hue sets were vertical saccades, which are hard to explain based on any lateral bias. There still remains the possibility that microstimulation at one site induced eye movement toward a particular direction for an unknown reason. To test this, we compared the observed bias in the direction of eye movements between stimulus sets. For example, in Figure 2c the animal frequently responded with a rightward saccade when microstimulation was applied with the green-saturation set, but with a leftward saccade with the blue-saturation and red-saturation sets. In the same manner, the animal responded with a downward saccade with the green-hue set, but with an upward saccade with the red-hue set. Similar inconsistency in the saccade direction was observed in cases that showed a large effect of microstimulation (Fig. 6d-f), and also for the entire population (data not shown). We therefore conclude that microstimulation of this subregion in the AIT modulated color signals.

### Spatial organization related to color in the AIT

Previous studies in unit recordings (Banno et al. 2010; Conway et al. 2007; Matsumora et al. 2008; Namima et al. 2014; Yasuda et al. 2010) and imaging methods (Conway et al. 2007; Conway and Tsao 2006; Harada et al. 2009; Tootell et al. 2004) have shown that there are subregions within the IT where color-selective activities are clustered. In the present study, we observed that within the effective hemisphere, the sites where clear modulation of color judgment behavior occurred were clustered, and their spatial extent was highly correlated with the distribution of color-selective cells. This suggests that there is a causal link between the activities of color-selective neurons in the AITC and color judgment behavior, and that this region consists of a functional domain specifically related to color judgment.

The spatial extent of the effect of microstimulation using our current range (20-50 μA) was presumably several hundred micrometers from the electrode tip (Histed et al. 2009; Ranck 1975; Stoney et al. 1968; Tehovnik et al. 2006). As we sampled the data at intervals of 2 mm, the observed spatial organization of the effect of microstimulation should be reliable.

A relatively large effect of microstimulation was observed for one hemisphere in both monkeys, but the dominant hemisphere was not consistent between the animals; it was the right hemisphere in monkey YU and left hemisphere in monkey RG. On the other hand, there was no clear laterality of the color-selective responses in the AITC in either of our previous studies (Koida and Komatsu 2007; Matsumora et al. 2008) or in the present study (Fig. 4e, f). One possible explanation is that there are differences in the way neurons with different color preferences are clustered. For example, the dominant hemisphere may have relatively strong clustering of neurons with specific color selectivity, while the other hemisphere does not. Another possible explanation is that even if the distribution of color-selective neurons is roughly the same across hemispheres, differences in neural processing in downstream areas exert a strong effect on color judgment through stimulation in the dominant hemisphere. A similar discrepancy between the selective neural responses and the effect of electrical stimulation was observed in the human fusiform gyrus (Rangarajan et al. 2014), where conscious face perception was evoked only by stimulation of the right hemisphere, though face-selective activities were observed in both hemispheres.

### Possible explanation for the direction of bias

It is generally thought that microstimulation-driven changes in neural activity are excitatory and are combined with visually driven activity in an additive manner (Clark et al. 2011). In the present study, however, the directions of the bias and the preferred color of neurons at the stimulation sites were roughly opposite, which appears to contradict those previous studies. One possible explanation is that the stimulation acted as if it provided an antagonistic surround against the sample stimulus. If microstimulation of the AITC evokes a color percept that extends over a large visual field, the color of the small sample stimulus (2 deg) in the central visual field may be contrasted with the evoked color, which would antagonistically modulate the perceived color of the sample stimulus. Consistent with this idea, microstimulation of the MT reportedly induces responses toward the opposite direction of the neurons when the stimulation sites preferred wide-field motion (Born et al. 2000). It may be that activating wide-field sites induces background motion, and antagonistic center-surround interactions induce an effect that is opposite of the center target. In a similar fashion, in the present study microstimulation in the AITC, where neurons have large receptive fields, might have evoked background color and induced modulation antagonistic to the center color. Indeed, electrical stimulation in the human medial fusiform cortex produced a subjective percept of color near the center of gaze, which was not localizable to a small region of the visual field (Murphey and Maunsell 2008).

Another possible cause of the opposite color effect is that microstimulation had an inhibitory effect on the neurons. Prolonged inhibition reportedly occurs after electrical stimulation of anesthetized animals (Berman et al. 1991; Butovas and Schwarz 2003; Chung and Ferster 1998), though direct measurement of the time course of the effect of microstimulation in the MT of alert macaques revealed a long-lasting excitatory effect on behavior (Masse and Cook 2010). In our experiment, microstimulation was applied at the same time as the visual stimulation. It therefore seems unlikely that an inhibitory aftereffect modulated color judgment. One could argue that the electrical stimulation would precede the visually driven neural responses, taking into account the neuron’s response latency, which is about 100 ms in IT (Baylis and Rolls 1987; Vogels and Orban 1994). However, microstimulation experiments in the IT have shown that biased judgment is towards the preferred stimulus category at the stimulation site, irrespective of when the microstimulation ends during the visual stimulus presentation period (Afraz et al. 2006). It is thus unlikely that an inhibitory effect of microstimulation can explain the opposite tendency between the direction of the shift and the preferred color indicated by the MUA.

Previous microstimulation studies in the IT examining causal links between face-selective and depth-selective neuron activities and perception showed an additive effect between microstimulation and visual stimulation (Afraz et al. 2006; Verhoef et al. 2012), but our results are at variance with those findings. So what accounts the difference between the present results and those of previous studies? Other than the difference in the visual attributes examined, there were two differences in the experimental procedures between the previous and present studies that may have affected the results. One difference is the stimulus size. Our color stimulus (2 deg in diameter) was smaller than the face stimulus (7 deg) in Afraz et al. and the depth stimulus (5 deg) in Verhoef et al. (Afraz et al. 2006; Verhoef et al. 2012). If the stimulation-induced percept is spatially spread, a contrast effect may become more prominent with smaller stimuli. The other difference is in the way discrimination difficulty was manipulated. Previous studies have used random dot noise to degrade face/non-face images (Afraz et al. 2006) or convex/concave stimuli (Verhoef et al. 2012), and the difficulty of the task was manipulated by the amount of noise. On the other hand, we did not use noise in the present study; instead, the difficulty was manipulated based on the distance between the ends of the spectrum, which made the range of stimulus variation very small. This may have made the contrast effect of the stimulation more visible in the present study for some unknown reason.

### Clustering of similarly tuned neurons

The modulatory effect of microstimulation on visually guided behavior is generally thought to reflect clustering of similarly tuned neurons. Although clustering of similarly color-tuned neurons has not been systematically studied in the AITC, the significant horizontal shift of the psychometric function seen in the present study suggests there is clustering of neurons that have similar color tunings. However, this does not necessarily mean that there is columnar organization of color tuning in the AITC. Previous studies examining the effects of microstimulation on motion speed in the MT (Liu and Newsome 2003) or motion direction in the MST (Celebrini and Newsome 1995) have shown that microstimulation modulates behavioral judgments, even though the columnar organization on these attributes is not obvious. Those authors assumed neurons with common preferred stimuli tended to cluster, whether or not the clusters actually formed columns.

The present study also suggests that, as a whole, the activated cells exhibit bias in color preference. If cells near the electrode tip have a wide variety on color preferences, and if the microstimulation does activate those cells, the evoked color signals would cancel each other, and the shift of color judgment behavior would be reduced. The actually observed shift in color judgment was significant and large, while the slope of the psychometric function did not consistently decrease, supporting the idea that there was population bias in color preference.

### Individual differences in directional bias

There was a difference in the observed directional bias between the two monkeys, which was distributed toward saturated color for monkey YU and along the line from blue for monkey RG (Fig. 3). This discrepancy might be due to the difference in the stimulus set. To examine this we conducted additional experiments with monkey RG using both the edge-color sets and the hue-and-saturation set (Fig. S3). In monkey RG, we did not observe the directional bias toward saturated colors observed in monkey YU. This suggests it is unlikely that the individual differences in the direction of bias are due to the difference in sample color sets. Taking into account the bias toward the direction opposite to the preferred color of neurons at the stimulation site (Fig. 7), the most parsimonious explanation is that the individual differences in directional bias reflects differences in the color preferences of the neuron populations studied between the two monkeys. That is, monkey YU had more neurons tuned to saturated colors, while monkey RG had more neurons tuned to blue or non-blue colors.

## Acknowledgment

We are grateful to M Togawa, M Takagi and T Ota for technical assistance. We are also grateful to T Matsumora for training monkeys. This study was supported by JSPS KAKENHI Grant Number, 19300113, 24300123, 22135007, and 15H05916 to HK and JSPS KAKENHI 25135718, and 15H05917 to KK. The authors declare no conflict of interest.

---

**Figure S1.**
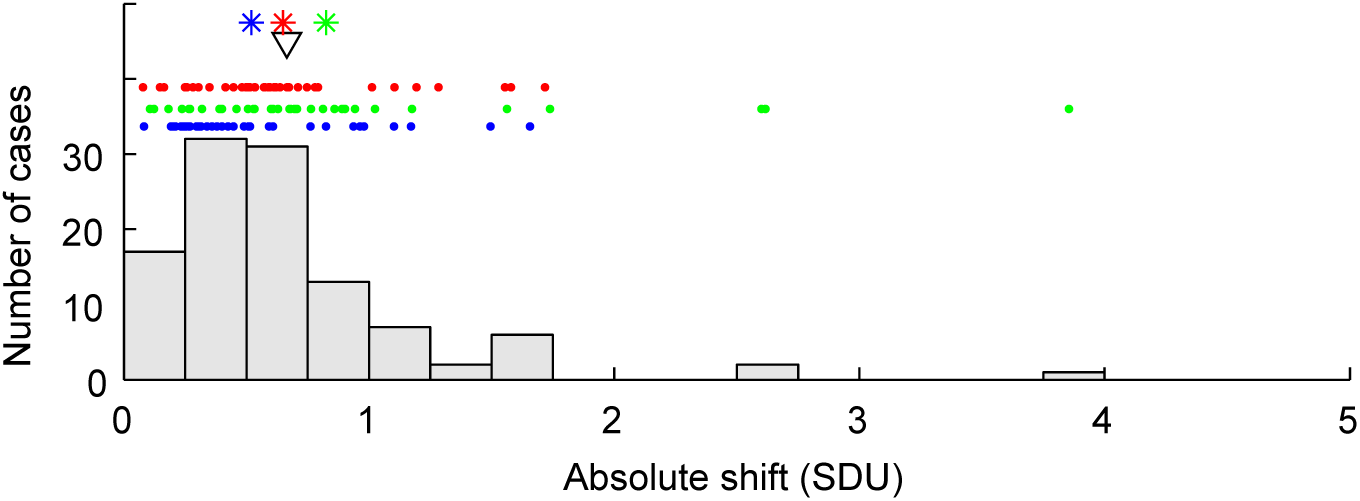
Magnitudes of the effects of microstimulation calculated as the root square sum of the effects with the hue and saturation sets in monkey RG (Fig. 3c) are illustrated in a histogram and raster plot. In the histogram, data from sets with different colors are combined. In the raster plot, data from each color set is shown separately in three colors. The triangle and asterisks indicate the mean shift.

**Figure S2.**
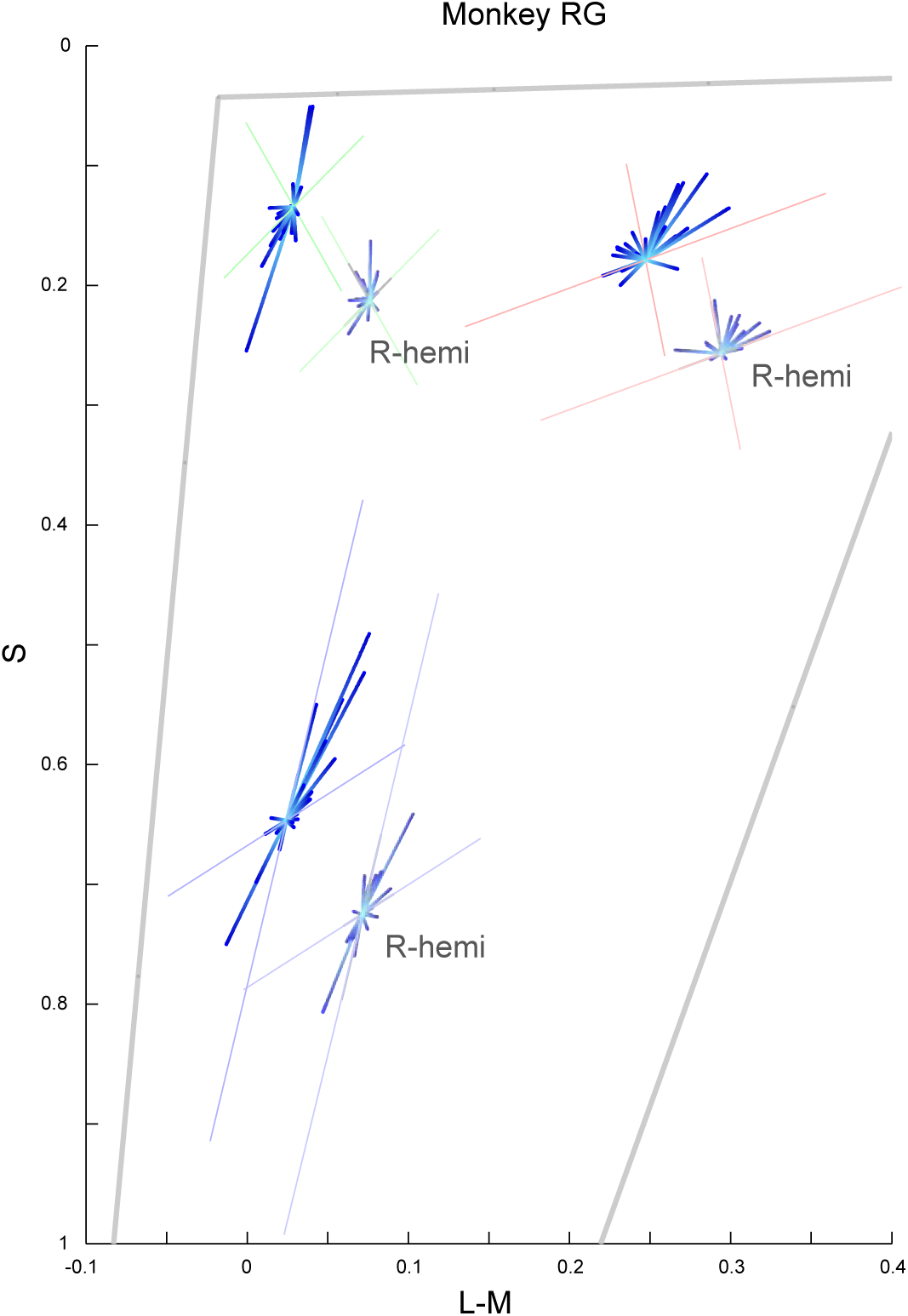
Direction of the effect of microstimulation plotted in cone space. In Figure 3d in the main text, the effect was plotted on CIE-xy chromaticity coordinates. However, a different color space – i.e., cone excitation space (Macleod-Boynton diagram, Macleod and Boynton, 1979) – might be useful for considering the relationship with early stage of color mechanisms such as cone output. Cone excitation was calculated using human cone fundamentals (Stockman et al., 1993). The power spectrum of the stimuli was measured with a spectrometer (PR-650, Photo Research, CA). The horizontal axis represents L-M cone output, while the vertical axis represents S cone output. The vertical axis is inverted to help compare the results shown in this diagram and those plotted on the CIE-xy chromaticity diagram. Consequently, blue is at the bottom, red is at the top right, and green is at the top left. Data obtained from the right hemisphere was slightly displaced for visualization purposes. The direction of the effect did not coincide with the S cone axis but was slightly rotated clockwise in this diagram.

**Figure S3.**
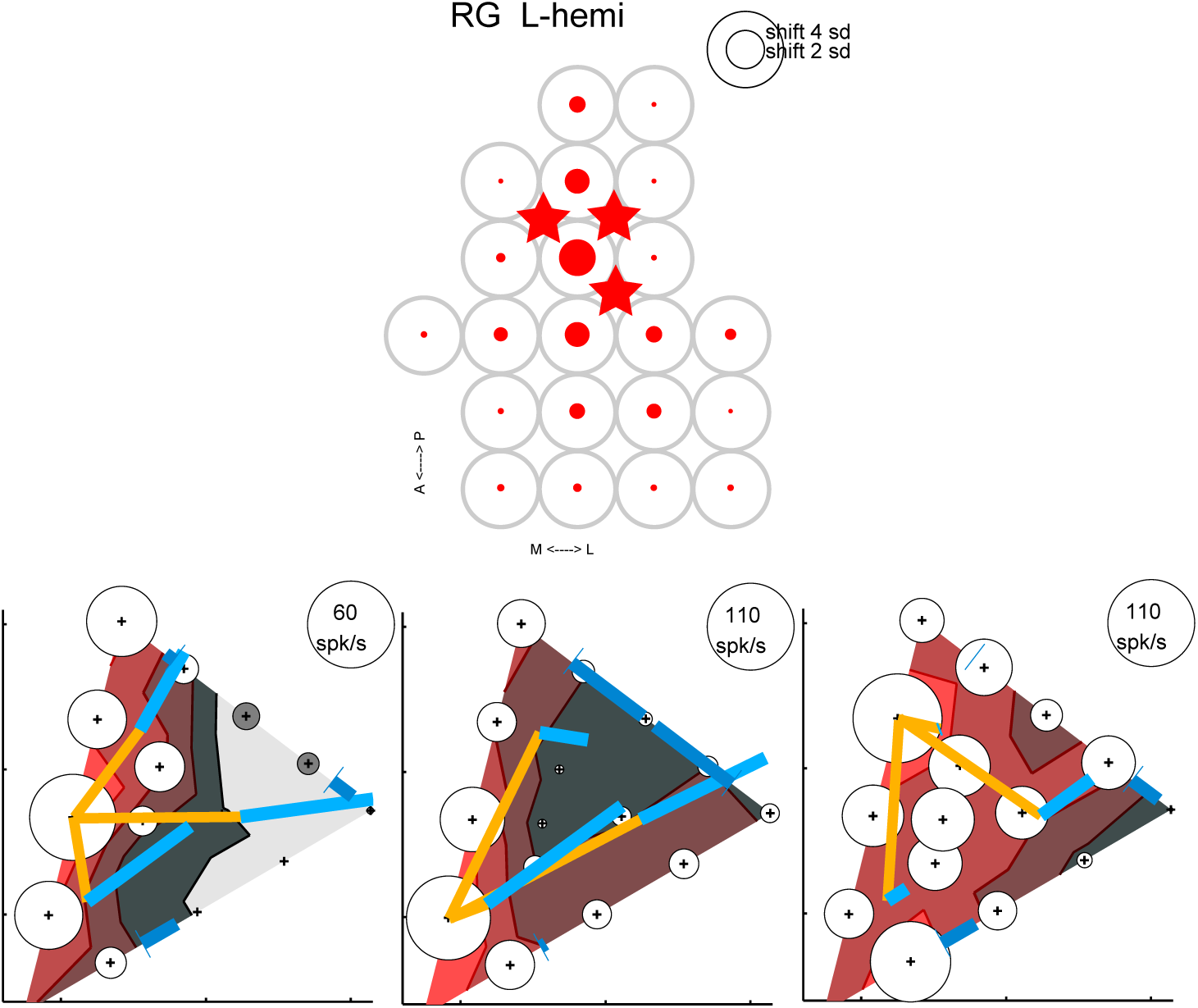
Results of an additional experiment with monkey RG using both edge-color sets and hue-saturation sets. In the main experiment, sample color sets differed between the animals. We found that the effects of microstimulation in monkey YU tended to direct toward more saturated colors, while in monkey RG they tended to distribute along a line from blue or toward blue (Fig. 3b, d). To test whether this discrepancy was due to a difference in color sample set used, we performed an additional experiment using the edge-color and hue-and-saturation color sets. This experiment was performed at three penetration sites in monkey RG, which are shown by stars in the upper panel. The results of the experiment are shown in the bottom row. The format is the same as in Figure 6 in the main text. Large effects of micro-stimulation were observed in all three cases, as were color-selective responses. The direction of the effects for edge-color sets were generally consistent with that of the hue-and-saturation color sets, and no clear bias in the direction of saturated colors was observed. The data for hue-saturation sets were included in the analysis in the main text

## References

Afraz SR, Kiani R, Esteky H. 2006. Microstimulation of inferotemporal cortex influences face categorization. Nature 442:692–695.

Banno T, Ichinohe N, Rockland KS, Komatsu H. 2010. Reciprocal Connectivity of Identified Color-Processing Modules in the Monkey Inferior Temporal Cortex. Cereb Cortex 21:1295–1310.

Baylis GC, Rolls ET. 1987. Responses of neurons in the inferior temporal cortex in short term and serial recognition memory tasks. Exp Brain Res 65:614–622.

Berman NJ, Douglas RJ, Martin KA, Whitteridge D. 1991. Mechanisms of inhibition in cat visual cortex. J Physiol 440:697–722.

Born RT, Groh JM, Zhao R, Lukasewycz SJ. 2000. Segregation of object and background motion in visual area MT: effects of microstimulation on eye movements. Neuron 26:725–734.

Buckley MJ, Gaffan D, Murray EA. 1997. Functional double dissociation between two inferior temporal cortical areas: perirhinal cortex versus middle temporal gyrus. J Neurophysiol 77:587–598.

Butovas S, Schwarz C. 2003. Spatiotemporal effects of microstimulation in rat neocortex: a parametric study using multielectrode recordings. J Neurophysiol 90:3024–3039.

Celebrini S, Newsome WT. 1995. Microstimulation of extrastriate area MST influences performance on a direction discrimination task. J Neurophysiol 73:437–448.

Chung S, Ferster D. 1998. Strength and orientation tuning of the thalamic input to simple cells revealed by electrically evoked cortical suppression. Neuron 20:1177–1189.

Clark KL, Armstrong KM, Moore T. 2011. Probing neural circuitry and function with electrical microstimulation. Proc Biol Sci 278:1121–1130.

Cohen MR, Newsome WT. 2004. What electrical microstimulation has revealed about the neural basis of cognition. Curr Opin Neurobiol 14:169–177.

Conway BR, Chatterjee S, Field GD, Horwitz GD, Johnson EN, Koida K, Mancuso K. 2010. Advances in color science: from retina to behavior. J Neurosci 30:14955–14963.

Conway BR, Moeller S, Tsao DY. 2007. Specialized color modules in macaque extrastriate cortex. Neuron 56:560–573.

Conway BR, Tsao DY. 2006. Color Architecture in Alert Macaque Cortex Revealed by fMRI. Cereb Cortex 16:1604–1613.

Crist CF, Yamasaki DS, Komatsu H, Wurtz RH. 1988. A grid system and a microsyringe for single cell recording. J Neurosci Methods 26:117–122.

Dean P. 1979. Visual cortex ablation and thresholds for successively presented stimuli in rhesus monkeys: II. Hue. Exp Brain Res 35:69–83.

DeAngelis GC, Cumming BG, Newsome WT. 1998. Cortical area MT and the perception of stereoscopic depth. Nature 394:677–680.

Desimone R, Albright TD, Gross CG, Bruce C. 1984. Stimulus-selective properties of inferior temporal neurons in the macaque. J Neurosci 4:2051–2062.

Desimone R, Schein SJ, Moran J, Ungerleider LG. 1985. Contour, color and shape analysis beyond the striate cortex. Vision Res 25:441–452.

Fujita I, Tanaka K, Ito M, Cheng K. 1992. Columns for visual features of objects in monkey inferotemporal cortex. Nature 360:343–346.

Harada T, Goda N, Ogawa T, Ito M, Toyoda H, Sadato N, Komatsu H. 2009. Distribution of colour-selective activity in the monkey inferior temporal cortex revealed by functional magnetic resonance imaging. Eur J Neurosci 30:1960–1970.

Heywood CA, Gaffan D, Cowey A. 1995. Cerebral achromatopsia in monkeys. Eur J Neurosci 7:1064–1073.

Histed MH, Bonin V, Reid RC. 2009. Direct activation of sparse, distributed populations of cortical neurons by electrical microstimulation. Neuron 63:508–522.

Horel JA. 1994. Retrieval of color and form during suppression of temporal cortex with cold. Behav Brain Res 65:165–172.

Huxlin KR, Saunders RC, Marchionini D, Pham HA, Merigan WH. 2000. Perceptual deficits after lesions of inferotemporal cortex in macaques. Cereb Cortex 10:671–683.

Judge SJ, Richmond BJ, Chu FC. 1980. Implantation of magnetic search coils for measurement of eye position: an improved method. Vision Res 20:535–538.

Koida K, Komatsu H. 2007. Effects of task demands on the responses of color-selective neurons in the inferior temporal cortex. Nat Neurosci 10:108–116.

Komatsu H. 1998. Mechanisms of central color vision. Curr Opin Neurobiol 8:503–508.

Komatsu H, Ideura Y. 1993. Relationships between color, shape, and pattern selectivities of neurons in the inferior temporal cortex of the monkey. J Neurophysiol 70:677–694.

Komatsu H, Ideura Y, Kaji S, Yamane S. 1992. Color selectivity of neurons in the inferior temporal cortex of the awake macaque monkey. J Neurosci 12:408–424.

Lafer-Sousa R, Conway BR. 2013. Parallel, multi-stage processing of colors, faces and shapes in macaque inferior temporal cortex. Nat Neurosci 16:1870–1878.

Liu J, Newsome WT. 2003. Functional organization of speed tuned neurons in visual area MT. J Neurophysiol 89:246–256.

Logothetis NK, Sheinberg DL. 1996. Visual object recognition. Annu Rev Neurosci 19:577–621.

Masse NY, Cook EP. 2010. Behavioral time course of microstimulation in cortical area MT. J Neurophysiol 103:334–345.

Matsumora T, Koida K, Komatsu H. 2008. Relationship between color discrimination and neural responses in the inferior temporal cortex of the monkey. J Neurophysiol 100:3361–3374.

Maunsell JH, Newsome WT. 1987. Visual processing in monkey extrastriate cortex. Annu Rev Neurosci 10:363–401.

Murphey DK, Maunsell JH. 2008. Electrical microstimulation thresholds for behavioral detection and saccades in monkey frontal eye fields. Proc Natl Acad Sci U S A 105:7315–7320.

Namima T, Yasuda M, Banno T, Okazawa G, Komatsu H. 2014. Effects of luminance contrast on the color selectivity of neurons in the macaque area v4 and inferior temporal cortex. J Neurosci 34:14934–14947.

Perrett DI, Rolls ET, Caan W. 1982. Visual neurones responsive to faces in the monkey temporal cortex. Exp Brain Res 47:329–342.

Ranck JB, Jr. 1975. Which elements are excited in electrical stimulation of mammalian central nervous system: a review. Brain Res 98:417–440.

Rangarajan V, Hermes D, Foster BL, Weiner KS, Jacques C, Grill-Spector K, Parvizi J. 2014. Electrical stimulation of the left and right human fusiform gyrus causes different effects in conscious face perception. J Neurosci 34:12828–12836.

Rolls ET. 1984. Neurons in the cortex of the temporal lobe and in the amygdala of the monkey with responses selective for faces. Hum Neurobiol 3:209–222.

Salzman CD, Britten KH, Newsome WT. 1990. Cortical microstimulation influences perceptual judgements of motion direction. Nature 346:174–177.

Salzman CD, Murasugi CM, Britten KH, Newsome WT. 1992. Microstimulation in visual area MT: effects on direction discrimination performance. J Neurosci 12:2331–2355.

Stoney SD, Jr., Thompson WD, Asanuma H. 1968. Excitation of pyramidal tract cells by intracortical microstimulation: effective extent of stimulating current. J Neurophysiol 31:659–669.

Takechi H, Onoe H, Shizuno H, Yoshikawa E, Sadato N, Tsukada H, Watanabe Y. 1997. Mapping of cortical areas involved in color vision in non-human primates. Neurosci Lett 230:17–20.

Tanaka K. 1996. Inferotemporal cortex and object vision. Annu Rev Neurosci 19:109–139.

Tehovnik EJ, Tolias AS, Sultan F, Slocum WM, Logothetis NK. 2006. Direct and indirect activation of cortical neurons by electrical microstimulation. J Neurophysiol 96:512–521.

Tootell RB, Nelissen K, Vanduffel W, Orban GA. 2004. Search for color ‘center(s)’ in macaque visual cortex. Cereb Cortex 14:353–363.

Verhoef BE, Vogels R, Janssen P. 2012. Inferotemporal cortex subserves three-dimensional structure categorization. Neuron 73:171–182.

Vogels R, Orban GA. 1994. Activity of inferior temporal neurons during orientation discrimination with successively presented gratings. J Neurophysiol 71:1428–1451.

Yasuda M, Banno T, Komatsu H. 2010. Color selectivity of neurons in the posterior inferior temporal cortex of the macaque monkey. Cereb Cortex 20:1630–1646.

Zeki S. 2005. The Ferrier Lecture 1995 behind the seen: the functional specialization of the brain in space and time. Philos Trans R Soc Lond B Biol Sci 360:1145–1183.

## References

MacLeod DI, Boynton RM. 1979. Chromaticity diagram showing cone excitation by stimuli of equal luminance. J Opt Soc Am 69:1183–1186.

Stockman A, MacLeod DI, Johnson NE. 1993. Spectral sensitivities of the human cones. J Opt Soc Am A 10:2491–2521

